# NANOCUTSIGHT: A NANOPORE-SEQUENCING APPROACH AND ANALYSIS PIPELINE TO ASSESS GENOME EDITING EFFICACY IN VARIOUS CELL POPULATIONS

**DOI:** 10.64898/2026.09.03.748616

**Authors:** Danny Bergeron, Véronique Gaudreault, Maxime Duval, Sonya Nassari, François Boudreau, Mathieu Durand, Karine Choquet, Steve Jean

## Abstract

Genome editing has revolutionized biomedical sciences and is now an essential tool to define molecular pathways through genetic interaction and loss-of-function studies. Through its diverse variations, it allows for the generation of specific knockout cell lines or organisms, as well as the creation of endogenously edited gene regions. While high-throughput methodologies exist to map CRISPR/Cas9 genetic modifications, the validation of guide efficiencies in cell populations is often performed through analysis of the targeted gene product by western blotting or by deconvolution of Sanger sequencing chromatograms using TIDE or ICE assays. Here, we highlight a rapid nanopore sequencing pipeline, which we have named NanoCutSight, to quantify the percentage of indels at a specific genomic locus and to identify the types of modifications generated. We also benchmarked the methodology on various guide RNAs and in both cultured cell and organoid models. We believe that NanoCutSight will simplify the analysis of complex sample editing and enable the rapid screening of edited samples.

## BACKGROUND

CRISPR/Cas methodologies are utilized worldwide to study gene function in various contexts (Li et al. 2023). The Cas9 enzyme is targeted to a specific genomic location by a guide RNA (gRNA) (Jackson et al. 2017), where it generates a double-strand break that is repaired either through Non-Homologous End Joining (NHEJ) or by homologous recombination (HR) (J. Y. Wang and Doudna 2023). NHEJ repair is imprecise and results in the generation of insertions or deletions (referred to as indels thereafter) at the targeted locus, which in most cases disrupts the open reading frame of the targeted region and blocks protein expression (Lin et al. 2014; Chakrabarti et al. 2019). While some studies have established algorithms to predict highly efficient gRNAs (Doench et al. 2016), their actual efficiencies can be influenced by numerous factors, and most predictions have not been experimentally validated. As such, it is essential to easily monitor the proportion and type of indels present in genetically modified cells, whether in clonal populations or in heterogeneous cell populations, to assess guide efficiency.

High-throughput methodologies have been developed to identify CRISPR edits in cells (Wienert et al. 2019; Tsai et al. 2015, 2018; Kim et al. 2015). These approaches rely on Illumina sequencing platforms and, while informative, have the drawbacks of high cost and often require a dedicated bioinformatician. To circumvent this, two straightforward methodologies have been established that rely on Sanger sequencing. Both *Tracking of Indels by Decomposition* (TIDE) and *Inference of CRISPR edits* (ICE) approaches use chromatogram deconvolution and allow the estimation of editing efficiency for a given gRNA (Brinkman et al. 2014; Conant et al. 2022). However, they do not provide an easy way to identify the different types of modifications present at a specific locus. This is particularly important because indels that are multiples of three should minimally affect the translation reading frame; therefore, precise characterization of indels within a population is essential. Additionally, highly validated antibodies are not available for most proteins (Ayoubi et al. 2023), further emphasizing the importance of accurately identifying CRISPR edits to predict their functional impact.

Nanopore long-read sequencing now offers the cost-effectiveness of Sanger sequencing (Cuber et al. 2023), with the additional advantage of enabling detailed analysis of thousands of sequence reads from a single run (Y. Wang et al. 2021). Thus, multiple PCR fragments can be analyzed simultaneously at low cost, with minimal bioinformatics requirements. Given these advantages, nanopore sequencing represents an appealing alternative to TIDE or ICE assays to map CRISPR/Cas9 genetic modifications in various contexts. The CRISPResso2 tool (Clement et al. 2019) was developed to analyze indels and insertions from CRISPR/Cas9 editing by Illumina sequencing, and has now been modified to accommodate nanopore sequencing (McFarlane, Polanco, and Bogema 2024). However, the algorithm was not adapted for key features of nanopore sequencing, such as noisy reads of variable lengths. In comparison, CRISPRNano, was developed as a suite to monitor various types of genetic modifications (i.e. homology driven recombination, prime editing and base editing) and to support Illumina, nanopore and nanopore/PacBio duplex reads. It also accounts in part for noisy reads from nanopore sequencing and allows for the analysis of multiple clones in parallel (Nguyen et al. 2022). Finally, GREPore-Seq (Quan et al. 2023) was developed to utilize nanopore sequencing data, but its analysis algorithm was focused to identify a wide array of insertions, from short double stranded oligonucleotides (dsODN) to large fragment insertions (plasmids), and not to map smaller indels.

To complement these tools and to support academic laboratories that often generate a more limited number of edited lines, we developed a simple and rapid analysis tool that enables efficient analysis of CRISPR edits across various cellular contexts. We named this pipeline NanoCutSight.

## MATERIAL AND METHODS

### Oligonucleotide synthesis and cloning

To benchmark the bioinformatic pipeline for the accurate identification of indels and insertions, we synthesized a 423 base pairs (bp) gBlock from the human gene *CCDC53* (IDT, Coralville, Iowa) along with identical fragments harboring a 1 bp insertion or deletion, or 5 bp and 50 bp deletions (Figure 1A and Supplementary Table 1). These different DNA fragments were mixed at various ratios to optimize NanoCutSight parameters. For cell- and organoid-based experiments, we used two previously validated gRNAs targeting the *FAM21C* (TCTCATTCCTAAAGTCCAGG) (Del Olmo et al. 2019) and *RAB14* (AGCGATTTAGGGCTGTTACA) (Gaudreault et al. 2025) genes and a highly efficient predicted *ATG16L1* gRNA (GAATAACCAAATGCAGCGGA). For all gRNAs, short complementary oligonucleotides with overlaps to BbsI were synthesized by IDT, annealed, and inserted into PX330A-1x2 (Addgene, 58766) digested with BbsI. All plasmids were validated by Sanger sequencing.

**Figure 1:**
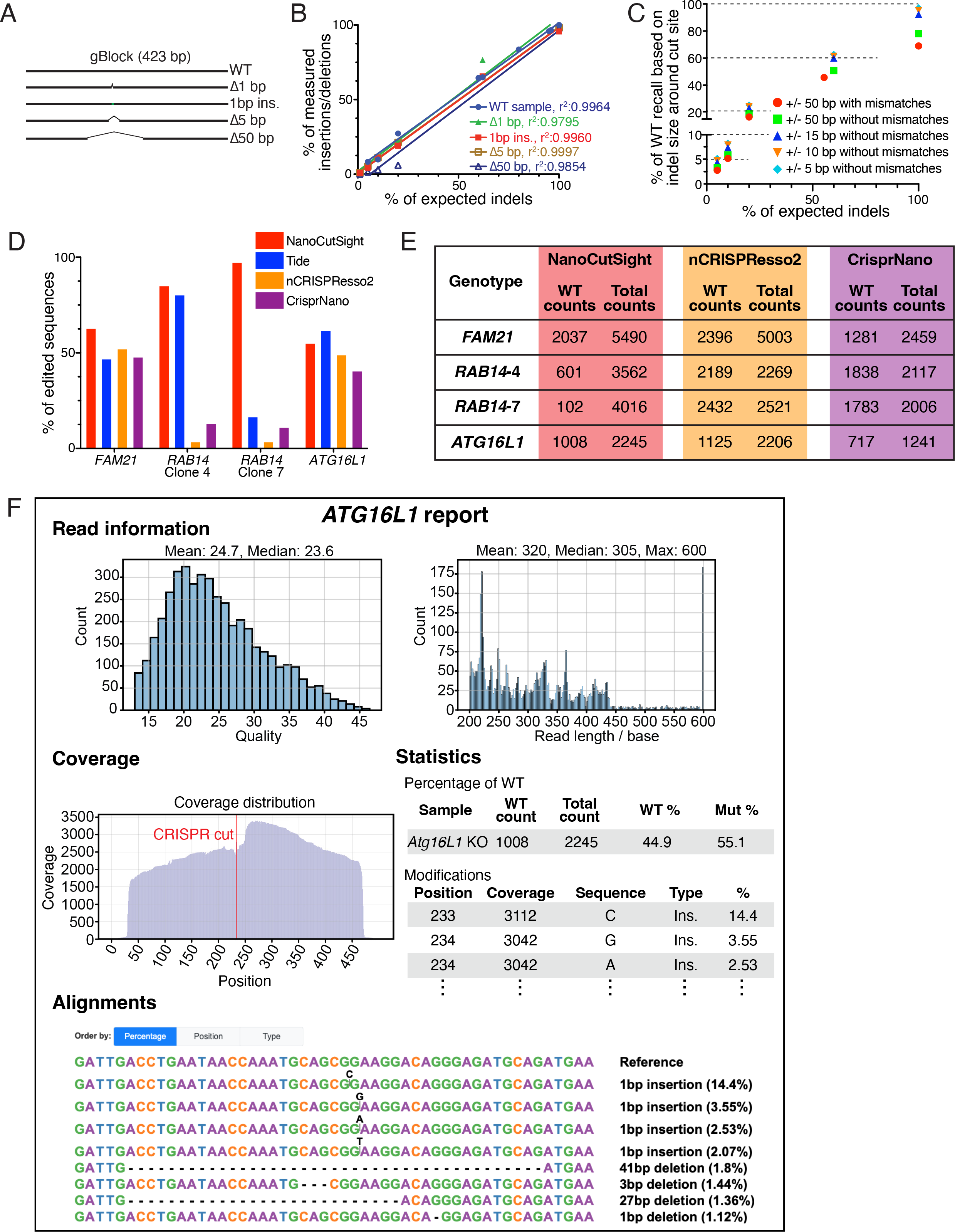
NanoCutSight enables the straightforward interpretation and analysis of genome-editing efficiency in various cellular contexts. (A) Cartoon representation of gBlocks used for NanoCutSight pipeline optimization. (B) Correlation analysis of the various gBlocks mixed with WT sequences at different ratios. The x-axis corresponds to the expected percentage of each modified gBlock added, while the y-axis corresponds to the measured percentage of modified gBlocks. (C) Influence of window size around the cut site and consideration of base substitutions on the recovery of wild-type gBlock sequences. (D) NanoCutSight, TIDE, CRISPResso2, and CRISPRNano show a comparable capacity to identify the percentage of modified indels in cell populations, but NanoCutSight performs more robustly in clonal cell lines. (E) Comparison of read depths for NanoCutSight, CRISPResso2, and CRISPRNano. **(F)** Representative overview of a report generated by NanoCutSight following gene-editing analysis of a cell population.

### Cell and organoid cultures

HeLa M cells were a kind gift from Pr. T. Yoshimori (University of Osaka). Cells were grown in Dulbecco’s Modified Eagle Medium (DMEM) (Wisent, St-Bruno, Qc) containing 5% fetal bovine serum and 1× penicillin/streptomycin (Wisent) at 37°C and 5% CO_2_.

Human colon organoids (ECO2) were obtained from Molecular Devices. Organoids were embedded in growth factor-reduced Matrigel and cultured in human colon organoid expansion media at 37°C in a humidified incubator with 5% CO2. Media was refreshed every 2 to 3 days.

### Transfection, DNA extraction and PCR

8 × 10⁵ HeLa M cells were seeded per well in a 6-well plate. The following day, the medium was removed and replaced with 2 ml of complete DMEM lacking antibiotics. Cells were transfected with a total of 1 µg of their respective gRNA plasmid and the pEGFP-Puro plasmid (Addgene; 45561) at a ratio of 15:1 using JetPrime (Polyplus, New York, NY, USA), following the manufacturer’s instructions. After a 6 h incubation, the medium containing the transfection mixture was replaced with complete DMEM for 24 h. Following this recovery period, transfected cells were selected for 36 h by the addition of 1 µg/ml puromycin. After the 36 h selection, cells were expanded for 72 h and lysed in QuickExtract DNA Extraction Solution (Lucigen, QE09050, Lystrup, Denmark), according to the manufacturer’s protocol. Five microliters of extracted genomic DNA were amplified with the Phusion High-Fidelity PCR Master Mix with HF buffer (New England Biolabs Ltd, M0531L, Whitby, ON, CA). For all PCRs, the gRNA binding region was centered, and PCR fragment lengths were 614 bp, 677 bp and 500 bp for *FAM21C*, *RAB14* and *ATG16L1*, respectively (Supplementary Table 1 for all sequences).

Human colorectal organoids were dissociated and co-electroporated with a peGFP-PURO and pspCas9-Atg16L1 sgRNA expressing plasmids, using NEPA21 electroporator (Nepagene). Electroporated organoids were then selected using puromycin treatment (2µg/ml for 72 h). Genomic DNA was prepared from organoids after 3 passages and amplified genomic DNA extraction and downstream analysis of CRISPR-induced modifications.

### TIDE analyses

For TIDE analyses, PCR amplicons from parental and edited HeLa M or human colon organoids were amplified and validated by gel electrophoresis. Following an in-column purification (New England Biolabs Ltd.; T1130S), PCR samples were sequenced by Sanger Sequencing at the Genomics Center at the CHUL Research Center (Université Laval), chromatograms (.abi files) were analyzed on the TIDE website (https://apps.datacurators.nl/tide/) and the percentage of indels were used to compare with NanoCutSight, CRISPResso2 and CRISPRNano.

### Nanopore sequencing and analysis

DNA from the gBlocks or from the PCR amplicons was purified using AMPure XP beads at a 1.8× ratio. Sequencing libraries were prepared from 400 ng of purified DNA with the Rapid Barcoding Kit (SQK-RBK114.94, Oxford Nanopore Technologies) following the manufacturer’s instructions and loaded onto a PromethION flow cell for sequencing. Raw signal data were base-called with Dorado using the super-accurate model for R10.4.1 chemistry and demultiplexed by barcode. Demultiplexed FASTQ files were used as input for downstream analysis.

### CRISPResso2 and CRISPNano analyses

CRISPResso2 was used via the https://crispresso2.pinellolab.org/submission link. Cas9 was selected as the editing tool, and the “single-end reads” sequencing option was chosen. Nanopore sequencing FASTQ files were provided as R1 input, along with the amplicon and sgRNA sequences. CRISPRNano was accessed through (www.crisprnano.de). Briefly, FASTQ files generated from nanopore sequencing were uploaded, and “nanopore” was selected as the data type. The maximum expected genotype number was set to 10, and “boosting mode” was not enabled. gRNA sequences were provided in the gRNA/pegRNA field. In the optional parameters, the amplicon sequence was entered under the “Custom Sequence” tab, 90 was specified in the “Interest Region” field, and an indel threshold of 2 was applied in the corresponding parameter.

## IMPLEMENTATION

Demultiplexed FASTQ files are processed with the NanoCutSight pipeline, a Snakemake (v7.24) workflow (available at https://github.com/dannyxbergeron/NanoCutSight. Reads from each bar-code are merged and filtered with seqkit (v2.4.0). 20 bp are trimmed from both ends, reads shorter than 200 bp or with a mean Phred quality below 13 are discarded, and reads are truncated to 1.2× the expected amplicon length. Because nanopore reads are obtained in both orientations, read strand is then corrected with restrander v1.1.3 (https://github.com/jakob-schuster/restrander) so that all reads align consistently to the reference strand and indel positions relative to the cut site are directly comparable across reads. Processed reads are aligned to a sample-specific reference sequence (provided as FASTA) with minimap2 (v2.26) in two passes. A splice-aware alignment (-x splice) is sorted and indexed with samtools (v1.15) and is used to compute per-position coverage and uniquely mapped reads, and a base-level alignment (-x map-ont --cs=long) generates long-form cs difference tags used for variant calling. Insertions and deletions are parsed from the cs tags and scored within a +/-10 bp window centered on the predicted Cas9 cut site. Only positions covered to >100x are considered, and indels present in 1% or greater of reads are retained. The editing efficiency is defined as the fraction of reads carrying an insertion or deletion overlapping the cut-site window, with base substitutions ignored. A per-position background base-calling error rate is estimated as the mean per-position frequency of mismatched bases (substitutions, insertions, and deletions) at positions outside the cut-site window. Finally, the pipeline assembles an interactive HTML report summarizing read-quality and read-length distributions, amplicon coverage, the list of detected indels with their frequencies, sequence alignments of the predominant variants, and the percentage of wild-type alleles.

## RESULTS

While Sanger sequencing-based chromatogram deconvolution by TIDE or ICE allows estimation of the percentage of modified cells (Brinkman et al. 2014; Conant et al. 2022), these approaches do not easily map the type and proportion of each modification. On the other hand, CRISPResso2 and CRISPRNano allows for the use of nanopore sequencing data, but they were not specifically designed for indel detection and thus have some limitations. We therefore focused on developing a nanopore sequencing–based approach enabling rapid analysis of indels from PCR amplicons. To initially test and benchmark the methodology and the analysis algorithm, we generated four independent gBlock fragments containing different deletion lengths or a single base pair insertion (Figure 1A and Supplementary Table 1). By combining different ratios of “mutated” gBlocks with wild-type fragments and subjecting them to nanopore sequencing, we observed a strong correlation (average R^2^=0.9914) between the expected and measured proportions of mutated gBlocks (Fig. 1B).

Importantly, and as reported by others (Quan et al. 2023), we noted that nanopore base-calling errors occurred even though all gBlock sequences had been validated (Tan et al. 2022). These errors were not uniformly distributed along the amplicon. Most positions were called accurately, whereas a subset of positions recurrently showed errors in a large fraction of reads, including homopolymer tracts and other contexts the base-calling model resolves less reliably. We quantified this background as the fraction of reads differing from the reference at each position, averaged over positions outside the cut-site window. The mean was ∼0.8% (median ∼0.7%), inflated by a tail of hotspot positions (approximately one in eighteen, reaching up to ∼7%), while the remaining ∼95% of positions had a median error of ∼0.6%. When pure mutated gBlocks were sequenced, 1.6–4.3% of reads were not assigned to their designed modification and were instead miscalled as wild-type, as an alternative indel class, or left unassigned. On real samples this back-ground can be further inflated by mismatches between the user-provided reference and the sample’s actual sequence (including allele-specific variants) and by base-calling errors in homopolymer tracts, and is therefore best interpreted as an upper bound on base-calling error. These observations confirm that nanopore base-calling errors can influence data interpretation and motivated restricting indel scoring to the +/-10 bp window around the cut site, where true CRISPR indels occur at frequencies far exceeding this background (Chakrabarti et al. 2019).

To assess the optimal indel analysis window and facilitate alignment generation, we evaluated the impact of varying the size of the analysis window around the predicted CRISPR/Cas9 cut site using generated gBlocks. The analysis was centered on the region surrounding the Cas9 cut site, as most indels are expected to occur proximal to this location (Chakrabarti et al. 2019). We also assessed if base-calling errors would affect the analysis and whether mismatches (from reverse transcriptase or PCR errors and from base-calling issues) should be included in the analysis. Comparing various analysis windows highlighted that including mismatches decreased the percentage of wild-type sequence recall, which was evident when wider deletions were considered (Fig. 1C, red dots compared to green squares). When mismatches were removed, we did not observe major differences between +/-5, +/-10 and +/-15 bp analysis windows for WT recall amongst various WT DNA gBlock percentages. Hence, to ease analysis and to ensure that most indels would be recovered, we opted to score indels within a 20 bp window (+/− 10 bp) around the targeted site; we also excluded base substitutions, focusing only on deletions and insertions.

To evaluate the methodology across different contexts, we generated PCR amplicons from HeLa cell populations targeted at the *FAM21C* gene using a previously validated gRNA (Del Olmo et al. 2019). All PCR amplicons were analyzed using NanoCutSight, TIDE, CRISPResso2, and CRISPRnano. We observed strong concordance between the methodologies in clonal populations (Fig. 1D). NanoCutSight estimated an indel frequency of 62.9% within the population, whereas TIDE provided an estimate of 47.9%. We further applied NanoCutSight to clonal *RAB14* knockout (KO) cells (Gaudreault et al. 2025) and were able to determine around 84% editing in clone #4 and close to a 100% editing in clone #7. Surprisingly, all other approaches, except for TIDE in clone KO-4, failed to identify these cells as clonal populations bearing specific indels (Fig. 1D). This discrepancy is unlikely to be due to insufficient read depth, as more than 2,000 reads were analysed by both CRISPResso2 and CRISPRNano (Fig. 1E). Rather, these differences may be attributable to variations in alignment algorithms, as these methods allow the mapping of a broader range of genetic modifications, whereas NanoCutSight specifically focuses on indels proximal to the cut site.

Finally, to evaluate the methodology in a different cellular context, we applied it to a gene-edited human colon organoid population at the *ATG16L1* locus. Consistent with observations in cell lines, NanoCutSight identified a broad range of indels (55%) (Fig. 1D). Notably, we detected nine independent indels representing more than 1% of total reads, along with multiple additional indels cumulatively accounting for a further 9% of modified cells. Given that each organoid derives from a single stem cell, these results suggest that NanoCutSight could be used to track the clonal expansion of distinct edited organoid populations. Altogether, these findings highlight the strength of this analytical pipeline in detecting indels across diverse cellular populations and enable detailed characterization of CRISPR/Cas9 editing across a range of experimental and biological contexts.

To further facilitate the use of NanoCutSight for analyzing CRISPR/Cas9-induced indels, we constructed a Snakemake pipeline to perform the analysis and to generate a validation report. The analysis report summarizes key information from nanopore sequencing runs (Fig. 1F). To run the pipeline, users provide a sample sheet containing, for each sample, the sequencing barcode, sample name, expected amplicon length, predicted CRISPR cut position, and expected wild-type sequence (FASTA format), along with a table pairing each edited sample with its corresponding wild-type control. From the corresponding demultiplexed FASTQ file, the pipeline generates a comprehensive report that includes quality-control metrics (histograms of read quality, read length distribution, and PCR fragment coverage), the percentage of wild-type alleles, a table of detected modifications along with their relative frequencies, and sequence alignments for the most predominant variants. This output enables users to readily assess and document the efficiency of a given gRNA at its target genomic locus.

## DISCUSSION

Here, we developed NanoCutSight to enable straightforward analysis of CRISPR/Cas9-mediated genome editing across diverse cellular contexts. NanoCutSight complements existing tools such as TIDE, ICE, CRISPResso2, and CRISPRNano by providing an alternative approach for analyzing a wide range of indels at the level of individual alleles in clonal cell lines, as well as quantifying editing frequencies within heterogeneous cell populations across various sample types. In addition, the generated report includes quality-control histograms that allow rapid assessment of sequencing data quality and overall experiment performance.

One limitation of nanopore sequencing is the frequency of base-calling errors (Tan et al. 2022), which increases the apparent percentage of edited cells because a substantial proportion of reads do not perfectly align with the theoretical wild-type sequence. We addressed this by focusing the analysis on the region immediately surrounding the predicted Cas9 cut site, since the most frequent alterations following CRISPR/Cas9 editing are 1 bp deletion or insertions flanking the cut site (Chakrabarti et al. 2019). It is worth noting, however, that any deletion or insertion overlapping the +/-10 bp window is still considered in the analysis, even if the event extends beyond the window boundary as observed in Figure 1F. We anticipate that improvements in nanopore base-calling will eventually allow expansion of the analysis window around the cut site to easily allow measurements of single base edited samples. While CRISPResso2 and CRISPRNano allow for the analysis of a wide range of modifications in numerous samples, their limitation is that they did not perform well for single clones, while NanoCutSight solely focuses on indels close to targeted sites and led to results agreeing with protein expression data (Gaudreault et al. 2025). Given the increasing accessibility and low cost of nanopore sequencing, NanoCutSight meets an important need by enabling academic laboratories to efficiently analyze genome-editing efficiencies across a wide range of samples.

## CONCLUSIONS

In conclusion, we present a simple analysis pipeline based on nanopore sequencing to monitor gene editing in various cell contexts. We envision that NanoCutSight will provide an effective alternative to TIDE, ICE, CRISPResso2 and CRISPRNano and enable improved characterization of CRISPR/Cas9-modified cells.

## AVAILABILITY AND REQUIREMENTS

**Project name:** NanoCutSight

**Project home page:** https://github.com/dannyxbergeron/NanoCutSight

**Operating system(s):** Linux (tested on Ubuntu)

**Programming language:** Python 3, Snakemake, Bash; HTML/JavaScript (Bootstrap, DataTables) for the report template

**Other requirements:** Snakemake ≥ 7, Conda/Mamba, a C++17 compiler and zlib development headers (used once, to compile restrander automatically on first run); all other dependencies are installed automatically as pinned conda environments.

**License:** GNU GPL v3

**Any restrictions to use by non-academics:** None

## DATA AVAILABILITY

The NanoCutSight source code, Snakemake workflow files, configuration files, documentation and amplicon reference sequences are freely available at https://github.com/dannyxbergeron/NanoCutSight under the GNU General Public License v3.0 or later.

## LIST OF ABBREVIATIONS

bp: Base pair(s)
Cas9: CRISPR-associated protein 9
CRISPR: Clustered Regularly Interspaced Short Palindromic Repeats
DMEM: Dulbecco’s Modified Eagle Medium
DNA: Deoxyribonucleic acid
dsODN: Double-stranded oligodeoxynucleotide
FASTA: Fast-All sequence format
GREPore-Seq: Genome Editing Read Processor for Oxford Nanopore Sequencing
gRNA: Guide RNA
HF: High Fidelity
HR: Homologous Recombination
ICE: Inference of CRISPR Edits
indel(s): Insertion(s) and deletion(s)
KO: Knockout
NHEJ: Non-Homologous End Joining
PCR: Polymerase Chain Reaction
pegRNA: Prime Editing guide RNA
RNA: Ribonucleic acid
sgRNA: Single guide RNA
Snakemake: Scalable workflow management system
TIDE: Tracking of Indels by Decomposition
WT: Wild Type

## DECLARATIONS

### Ethics approval and consent to participate

Not applicable.

### Consent for publication

Not applicable.

### Availability of data and materials

The NanoCutSight Snakemake pipeline and example configuration files are available on GitHub at https://github.com/dannyxbergeron/NanoCutSight under the GNU General Public License v3. The exact version analysed in this study will be archived on Zenodo with a citable DOI upon acceptance.

### Authors’ contributions

DB, MD, SJ, and KC conceptualized the study and developed the idea of creating NanoCutSight. MD, VG and SN performed the experiments. DB developed and optimized the analysis pipeline with suggestions from VG, SN, MD, SJ and KC. FB obtained funding and assisted with experiments linked to organoids. SJ secured the funding for this work. All authors read and approved the final version of the manuscript.

## Acknowledgements

We would like to thank members of the Jean and Choquet labs and the RNomique core facility for discussion and suggestions throughout this work. The RNomique core is operationnally supported by the Faculté de médecine et des sciences de la santé. The Canadian National Organoid Network (CNON), funded by the Weston Family Foundation, supported the organoid experiments. SJ is a member of the FRQS-funded Centre de Recherche du CHUS and the Institut de Recherche sur le Cancer de l’Université de Sherbrooke and KC is a member of the FRQS-funded Centre de Recherche sur le Vieillissement. SJ is also a member of PROTEO, the regroupement québécois de recherche sur la fonction, l’ingénierie et les applications des protéines.

## REFERENCES

Ayoubi, Riham, Joel Ryan, Michael S. Biddle, Walaa Alshafie, Maryam Fotouhi, Sara Gonzalez Bolivar, Vera Ruiz Moleon, et al. 2023. “Scaling of an Antibody Validation Procedure Enables Quantification of Antibody Performance in Major Research Applications.” eLife 12 (RP91645): RP91645.

Brinkman, Eva K., Tao Chen, Mario Amendola, and Bas van Steensel. 2014. “Easy Quantitative Assessment of Genome Editing by Sequence Trace Decomposition.” Nucleic Acids Research 42 (22): e168.

Chakrabarti, Anob M., Tristan Henser-Brownhill, Josep Monserrat, Anna R. Poetsch, Nicholas M. Luscombe, and Paola Scaffidi. 2019. “Target-Specific Precision of CRISPR-Mediated Genome Editing.” Molecular Cell 73 (4): 699–713.e6.

Clement, Kendell, Holly Rees, Matthew C. Canver, Jason M. Gehrke, Rick Farouni, Jonathan Y. Hsu, Mitchel A. Cole, et al. 2019. “CRISPResso2 Provides Accurate and Rapid Genome Editing Sequence Analysis.” Nature Biotechnology 37 (3): 224–26.

Conant, David, Tim Hsiau, Nicholas Rossi, Jennifer Oki, Travis Maures, Kelsey Waite, Joyce Yang, et al. 2022. “Inference of CRISPR Edits from Sanger Trace Data.” The CRISPR Journal 5 (1): 123–30.

Cuber, Piotr, Darren Chooneea, Clementine Geeves, Silvia Salatino, Thomas J. Creedy, Claire Griffin, Laura Sivess, Ian Barnes, Ben Price, and Raju Misra. 2023. “Comparing the Accuracy and Efficiency of Third Generation Sequencing Technologies, Oxford Nanopore Technologies, and Pacific Biosciences, for DNA Barcode Sequencing Applications.” Ecological Genetics and Genomics 28 (100181): 100181.

Del Olmo, Tomas, Annie Lauzier, Caroline Normandin, Raphaëlle Larcher, Mia Lecours, Dominique Jean, Louis Lessard, Florian Steinberg, François-Michel Boisvert, and Steve Jean. 2019. “APEX2-Mediated RAB Proximity Labeling Identifies a Role for RAB21 in Clathrin-Independent Cargo Sorting.” EMBO Reports 20 (2): e47192.

Doench, John G., Nicolo Fusi, Meagan Sullender, Mudra Hegde, Emma W. Vaimberg, Katherine F. Donovan, Ian Smith, et al. 2016. “Optimized sgRNA Design to Maximize Activity and Minimize Off-Target Effects of CRISPR-Cas9.” Nature Biotechnology 34 (2): 184–91.

Gaudreault, Véronique, Benoit Marchand, Raphaëlle Larcher, Sonya Nassari, Francis Bourassa, Mathilde Moreau, Dominique Jean, François-Michel Boisvert, Marie A. Brunet, and Steve Jean. 2025. “A Proximity Map of RAB GTPases Delineates Roles for RAB14 in EARP Complex and UHRF1BP1 Endosomal Recruitments.” Communications Biology 8 (1): 1717.

Jackson, Simon A., Rebecca E. McKenzie, Robert D. Fagerlund, Sebastian N. Kieper, Peter C. Fineran, and Stan J. J. Brouns. 2017. “CRISPR-Cas: Adapting to Change.” Science 356 (6333): eaal5056.

Kim, Daesik, Sangsu Bae, Jeongbin Park, Eunji Kim, Seokjoong Kim, Hye Ryeong Yu, Jinha Hwang, Jong-Il Kim, and Jin-Soo Kim. 2015. “Digenome-Seq: Genome-Wide Profiling of CRISPR-Cas9 off-Target Effects in Human Cells.” Nature Methods 12 (3): 237–43, 1 p following 243.

Li, Tianxiang, Yanyan Yang, Hongzhao Qi, Weigang Cui, Lin Zhang, Xiuxiu Fu, Xiangqin He, Meixin Liu, Pei-Feng Li, and Tao Yu. 2023. “CRISPR/Cas9 Therapeutics: Progress and Prospects.” Signal Transduction and Targeted Therapy 8 (1): 36.

Lin, Steven, Brett Staahl, Ravi K. Alla, and Jennifer A. Doudna. 2014. “Enhanced Homology-Directed Human Genome Engineering by Controlled Timing of CRISPR/Cas9 Delivery.” eLife 3.

McFarlane, Gus Rowan, Jenin Victor Cortez Polanco, and Daniel Bogema. 2024. “CRISPR-Cas Guide RNA Indel Analysis Using CRISPResso2 with Nanopore Sequencing Data.” BMC Research Notes 17 (1): 205.

Nguyen, Thach, Haribaskar Ramachandran, Soraia Martins, Jean Krutmann, and Andrea Rossi. 2022. “Identification of Genome Edited Cells Using CRISPRnano.” Nucleic Acids Research 50 (W1): W199–203.

Quan, Zi-Jun, Si-Ang Li, Zhi-Xue Yang, Juan-Juan Zhao, Guo-Hua Li, Feng Zhang, Wei Wen, Tao Cheng, and Xiao-Bing Zhang. 2023. “GREPore-Seq: A Robust Workflow to Detect Changes after Gene Editing through Long-Range PCR and Nanopore Sequencing.” *Genomics*, Proteomics & Bioinformatics 21 (6): 1221–36.

Tan, Kar-Tong, Michael K. Slevin, Matthew Meyerson, and Heng Li. 2022. “Identifying and Correcting Repeat-Calling Errors in Nanopore Sequencing of Telomeres.” Genome Biology 23 (1): 180.

Tsai, Shengdar Q., Nhu T. Nguyen, Jose Malagon-Lopez, Ved V. Topkar, Martin J. Aryee, and J. Keith Joung. 2018. “Erratum: Corrigendum: CIRCLE-Seq: A Highly Sensitive in Vitro Screen for Genome-Wide CRISPR–Cas9 Nuclease off-Targets.” Nature Methods 15 (5): 394–394.

Tsai, Shengdar Q., Zongli Zheng, Nhu T. Nguyen, Matthew Liebers, Ved V. Topkar, Vishal Thapar, Nicolas Wyvekens, et al. 2015. “GUIDE-Seq Enables Genome-Wide Profiling of off-Target Cleavage by CRISPR-Cas Nucleases.” Nature Biotechnology 33 (2): 187–98.

Wang, Joy Y., and Jennifer A. Doudna. 2023. “CRISPR Technology: A Decade of Genome Editing Is Only the Beginning.” Science 379 (6629): eadd8643.

Wang, Yunhao, Yue Zhao, Audrey Bollas, Yuru Wang, and Kin Fai Au. 2021. “Nanopore Sequencing Technology, Bioinformatics and Applications.” Nature Biotechnology 39 (11): 1348–65.

Wienert, Beeke, Stacia K. Wyman, Christopher D. Richardson, Charles D. Yeh, Pinar Akcakaya, Michelle J. Porritt, Michaela Morlock, et al. 2019. “Unbiased Detection of CRISPR Off-Targets in Vivo Using DISCOVER-Seq.” Science (New York, N.Y.) 364 (6437): 286–89.

